# COL: a pipeline for identifying putatively functional back-splicing

**DOI:** 10.1101/2023.11.08.566217

**Authors:** Zheng Li, Bandhan Sarker, Fengyu Zhao, Tianjiao Zhou, Jianzhi Zhang, Chuan Xu

## Abstract

Circular RNAs (circRNAs) are a class of generally non-coding RNAs produced by back-splicing. Although the vast majority of circRNAs are likely to be products of splicing error and thereby confer no benefits to organisms, a small number of circRNAs have been found to be functional. Identifying other functional circRNAs from the sea of mostly non-functional circRNAs is an important but difficult task. Because available experimental methods for this purpose are of low throughput or versality and existing computational methods have limited reliability or applicability, new methods are needed. We hypothesize that functional back-splicing events that generate functional circRNAs (i) exhibit substantially higher back-splicing rates than expected from the total splicing amounts, (ii) have conserved splicing motifs, and (iii) show unusually high back-splicing levels. We confirm these features in back-splicing shared among human, macaque, and mouse, which should enrich functional back-splicing. Integrating the three features, we design a computational pipeline named COL for identifying putatively functional back-splicing. Different from the methods that require multiple samples, COL can predict functional back-splicing using a single sample. Under the same data requirement, COL has a lower false positive rate than that of the commonly used method that is based on the back-splicing level alone. We conclude that COL is an efficient and versatile method for rapid identification of putatively functional back-splicing and circRNAs that can be experimentally validated. COL is available at https://github.com/XuLabSJTU/COL.

## INTRODUCTION

Circular RNAs (circRNAs) are a class of endogenous, single-stranded, covalently closed, mostly non-coding RNAs that are produced by back-splicing that links a downstream splice-donor site with an upstream splice-acceptor site [1, 2]. Over one half of human protein-coding genes produce circRNAs [3] and millions of circRNAs have been identified in animals [4] and human cancer cells [5, 6]. It is reported that circRNAs have important regulatory functions, such as acting as miRNA [7, 8] or protein [9] sponge to regulate gene expression, acting as scaffolds to mediate the formation of complexes [10], and being translated into small functional peptides [11, 12]. Additionally, a few circRNAs have been implicated in cardiovascular disease [13], Alzheimer’s disease [14], cancer [6, 15, 16], and other diseases [17–19]. Notwithstanding, of millions of known circRNAs, those with demonstrated functions are a tiny fraction.

Our recent study of back-splicing in human, macaque, and mouse estimated that over 97% of back-splicing results from splicing error, supporting the error hypothesis of the origin of circRNAs and potentially explaining why only a tiny fraction of circRNAs have been found functional [20]. However, given the huge pool of circRNAs, it is reasonable to assume that not all functional circRNAs have been discovered. Identifying functional circRNAs is challenging because of the presence of orders of magnitude more nonfunctional circRNAs. One commonly used approach, which may be called the differential expression method (DEM) [21–23], detects differentially expressed circRNAs between two groups of samples (*e.g.*, patients vs. controls) and considers them potentially functional. This method is obviously inapplicable to a single sample or non-comparable samples.

Another commonly used approach may be named the expression level method (ELM) [8, 24, 25]. It treats highly expressed circRNAs as putatively functional under the assumption that a circRNA needs to have a sufficiently high expression to exert its function. While functional circRNAs are likely to be highly expressed, highly expressed circRNAs may not be functional because their high expressions may simply arise from the high expressions of their parental genes. Hence, ELM is expected to generate many false positives when used alone for discovering functional circRNAs.

The conservation of circRNAs across species is another useful criterion for identifying functional circRNAs (referred to as the conservation method or CM), because random splicing errors are unlikely to be conserved across species [26–28]. However, CM requires not only the sequenced genomes but also circRNA data from multiple species, hindering its wide application in non-model organisms.

In addition to the above three computational approaches, a high-throughput experimental screening method based on CRISPR-Cas13 was recently developed [29]. This experimental method improves the discovery of functional circRNAs, but it is complicated and costly and cannot be used in species without a CRISPR-Cas13 screening system. Therefore, there is a need for a simple, inexpensive, and widely applicable method for identifying putatively functional circRNAs. In this work, we propose and validate such a method.

## RESULTS

### Three features of functional back-splicing

If a circRNA is functional, the back-splicing that generates the circRNA should be considered functional back-splicing. Therefore, to identify functional circRNAs is to discover functional back-splicing. Functional back-splicing is expected to have the following three features. First, the error hypothesis of the origin of circRNAs asserts that most back-splicing results from deleterious splicing error, predicting a negative correlation between the amount of splicing at a splicing junction and the back-splicing rate at the junction, which was indeed observed [20]. Functional back-splicing is expected to break this rule and show a higher back-splicing rate than predicted by the splicing amount. Second, functional back-splicing is expected to have conserved splicing motifs. Third, functional back-splicing should have a relatively high back-splicing level to allow the production of sufficient circRNA molecules.

To empirically verify the above suggested characteristics of functional back-splicing, we should ideally compare functional back-splicing with the rest of back-splicing. However, because only a few instances of back-splicing are demonstrably functional, we instead compared human back-splicing that is shared with macaque and mouse with the rest of human back-splicing, because shared back-splicing is more likely than unshared back-splicing to be functional. Specifically, we first identified linearly spliced reads and back-spliced reads in appropriate RNA sequencing (RNA-seq) data from human, macaque, and mouse (see **Methods**). To reduce the impact of potential sequencing errors, we required at least two spliced reads to call a splicing event. We defined the total splicing amount at a splicing junction by the total number of spliced reads present in its donor and acceptor. Then, the back-splicing rate at a back-splicing junction was computed by dividing the number of back-splicing reads by the total splicing amount at the junction (see **Methods**). To ensure the reliability in the discovery of shared back-splicing, we defined back-splicing as shared if it has at least two back-spliced reads in the same tissue in each of human, macaque, and mouse (see **Methods**); all other human back-splicing was regarded as unshared.

We combined 11 human tissues to represent human and analyzed all the back-splicing. Consistent with our previous finding [20], a significant negative correlation exists between the back-splicing rate and the total splicing amount across splicing junctions (**Fig. 1A**). To identify back-splicing violating this general rule, we regressed the back-splicing rate against the splicing amount (**Fig. 1A**) and calculated Cook’s distance for each back-splicing event (see **Methods**). A back-splicing event is considered an outlier if its Cook’s distance exceeds four times the mean Cook’s distance of all back-splicing events (see **Methods**). We respectively observed 3,587 outliers above and 236 outliers below the regression line (**Fig. 1A**). The error hypothesis suggests that the outliers above the regression line are beneficial (i.e., functional) whereas those below the regression line are particularly deleterious. Indeed, shared back-splicing contains significantly more outliers above the regression line than does unshared back-splicing (*P* < 10^−16^, Fisher’s exact test; **Fig. 1B**). By contrast, shared back-splicing contains fewer outliers below the regression line than does unshared back-splicing (**Fig. S1**).

**Fig. 1.**
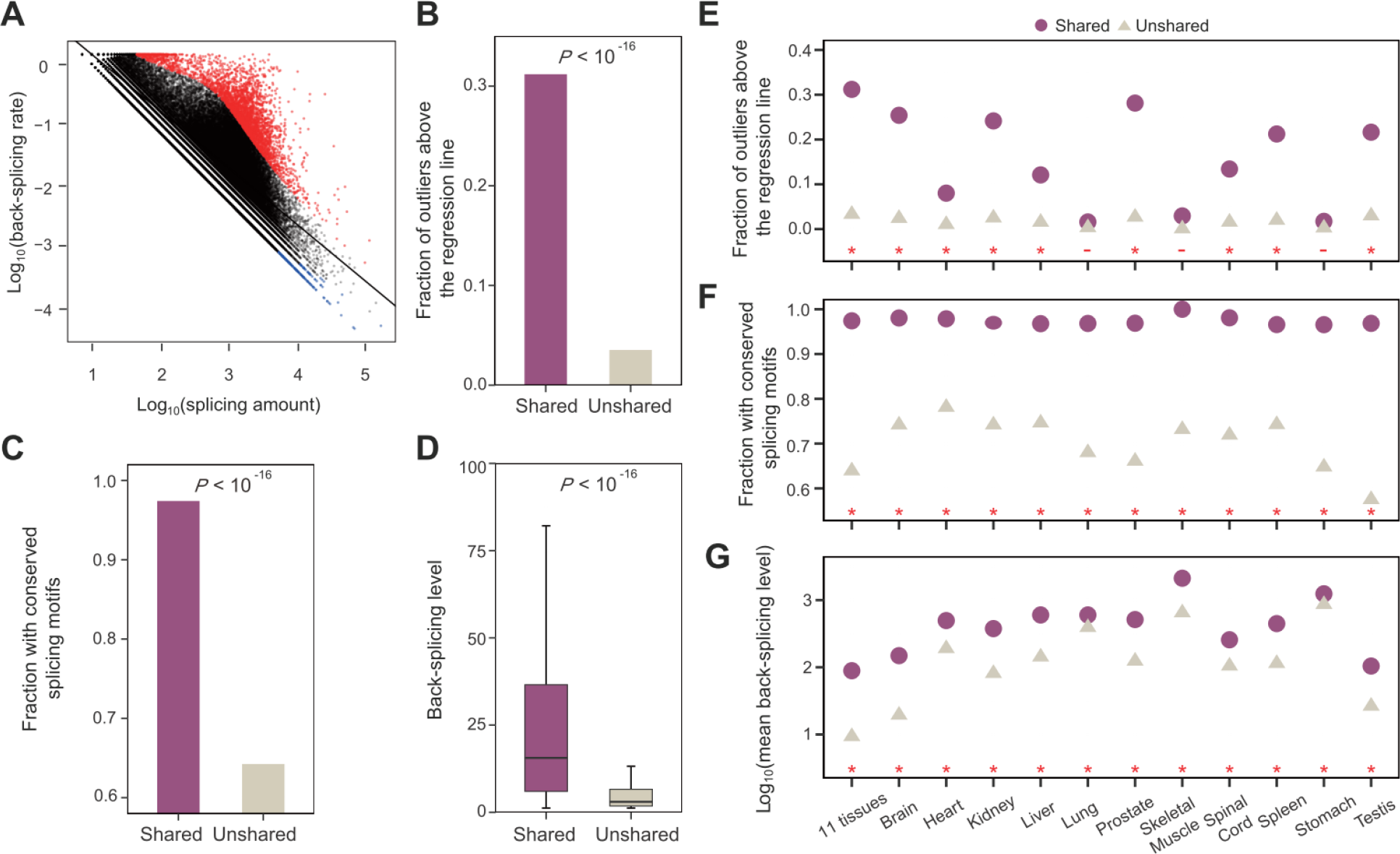
Comparison of three features between shared and unshared back-splicing in human tissues. (**A**) Linear regression between the back-splicing rate and splicing amount across back-splicing junctions. Each dot represents a back-splicing junction, and the solid black line shows the linear least-squares regression. Red and blue dots are outliers above and below the regression line, respectively. (**B**) Outliers above the regression line are enriched in shared back-splicing relative to unshared back-splicing. (**C**) Shared back-splicing has a higher fraction of conserved splicing motifs than does unshared back-splicing. (**D**) The back-splicing level is higher for shared than unshared back-splicing. Data in (A) to (D) are from the combination of 11 human tissues. (**E-G)** Comparison of the three features between shared and unshared back-splicing in each of 11 human tissues. Statistical significance of a difference is indicated by a dash (non-significant) or star (significant at *P* < 0.05) at the bottom of each panel. In (D) and (G), the back-splicing level at a back-splicing junction is measured by the number of back-spliced reads at the junction per million total back-spliced reads in the sample (BSRPM).

Both linear-splicing and back-splicing use GU/AG splicing motifs. We found that the splicing motifs of 97.4% of shared back-splicing are conserved across human, macaque, and mouse, significantly higher than the corresponding value (64.2%) for unshared back-splicing (*P* < 10^−16^, Fisher’s exact test; **Fig. 1C**).

We then measured the relative back-splicing level at a splicing junction by the number of corresponding back-splicing reads per million total back-splicing reads in the sample (BSRPM) [30, 31]. Indeed, BSRPM is higher for shared than unshared back-splicing (*P* < 10^−16^, Mann– Whitney U test; **Fig. 1D**).

Therefore, all three hypothesized features of functional back-splicing are verified in the shared back-splicing of the human tissues combined. We repeated the above analyses in each of 11 tissues in the dataset and observed qualitatively similar results (**Fig. 1E-G**).

### A feature-based integrative pipeline for predicting functional back-splicing

To examine the utility of the above three features for predicting shared (as a proxy for functional) back-splicing, we measured the level of enrichment of shared back-splicing in the candidate back-splicing identified using these features. In human, 2,318 (2.8%) of 83,458 back-splicing events are shared among human, macaque, and mouse. So, 2.8% is the baseline in our enrichment calculation. Of the 3,587 outliers over the regression line in **Fig. 1A**, 723 (or 20.2%) belong to shared back-splicing, a 7.2-fold enrichment relative to the baseline (*P* < 10^−16^, Fisher’s exact test; **Fig. 2A**). Similarly, 4.2% of back-splicing with conserved splicing motifs belong to shared back-splicing, a 1.5-fold enrichment relative to the baseline (*P* < 10^−16^, Fisher’s exact test; **Fig. 2A**). To compare the utility of back-splicing levels with that of regression outliers, we examined the top 3,587 back-splicing events in terms of their back-splicing levels and found that 22.5% of them belong to shared back-splicing, an 8.0-fold enrichment over the baseline (*P* < 10^−16^, Fisher’s exact test; **Fig. 2A**). Therefore, each of the three features alone has some power in predicting shared back-splicing.

**Fig. 2.**
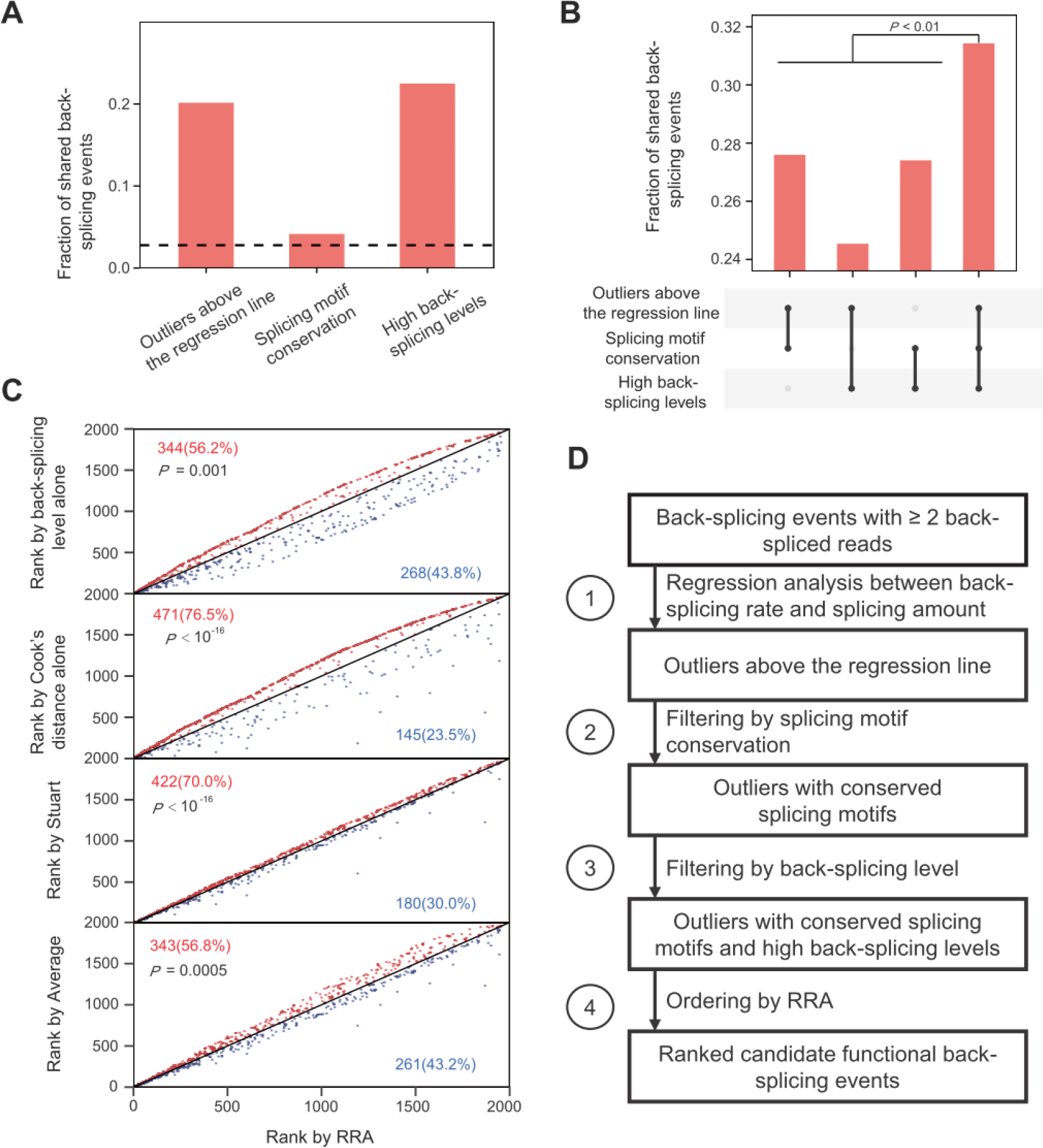
Predicting shared back-splicing using combinations of the three features. (**A**) The predictive power of each feature. The dotted line represents the baseline (i.e., the fraction of shared back-splicing events in all back-splicing events). (**B**) The predictive power of combined features through Venn diagram integration. *P*-value is from chi-squared test. (**C**) Pairwise comparison between RRA ranking and four other rankings. Each dot represents a back-splicing event. Dots above and below the diagonal are respectively colored in red and blue, with their numbers and fractions indicated. *P*-values are from binomial tests. (**D**) The integrative pipeline of COL for predicting functional back-splicing.

We next examined whether a combination of the three features has an enhanced power in predicting shared back-splicing. We first used the Venn diagram to integrate the back-splicing events identified by each feature (see **Methods**). We found that any combination of two features has a higher predictive power than any single feature (maximum *P* < 0.06, chi-squared test; **Fig. 2B**). The combination of all three features has the highest power—31.4% of the 1960 back-splicing events identified are shared (**Fig. 2B**).

While the above combination yields a set of candidates for shared back-splicing, it does not rank these candidates by their probabilities of sharing. To prioritize these candidates, we tested a few ordering methods such as Average, Stuart, and Robust Rank Aggregation (RRA) that can integrate different ranking results [32] (see **Methods**). Because the feature of splicing motif conservation is binary (i.e., conserved or not), we only considered Cook’s distance and back-splicing level in the ordering. In the pairwise comparison between any two ranking methods, we found the RRA ranking method to assign smaller rankings to a greater number of shared back-splicing events than any other ranking method examined (**Fig. 2C**, **Fig. S2**). Therefore, the RRA ranking method performs the best in prioritizing the candidates for shared back-splicing.

Based on the above results from the analysis of shared back-splicing as a proxy for functional back-splicing, we developed an integrative pipeline to predict functional back-splicing (**Fig. 2D**). It first identifies outliers above the regression line between the back-splicing rate and splicing amount across splicing junctions, then filters out those outliers without conserved splicing motifs, further filters out those without sufficiently high back-splicing levels, and finally prioritizes the candidates using RRA. Because this pipeline integrates <u>c</u>onservation of splicing motifs, <u>o</u>utliers in the regression analysis, and high back-splicing level, we refer to it as the COL method hereinafter. To facilitate the utilization of COL, we have developed an R package, which is available at https://github.com/XuLabSJTU/COL.

### Performance of COL

While the COL developed above can enrich shared back-splicing, how the predictive power of COL varies with the threshold of each feature considered is unclear. Therefore, we evaluated COL’s performance under different thresholds of the three features. As shown in **Fig. 2B**, the predictive power of COL (i.e., the fraction of identified back-splicing that is shared among human, macaque, and mouse) is 31.4% under the thresholds of (i) four times the mean Cook’s distance for the outliers, (ii) motif conservation between human and mouse, and (iii) a minimum back-splicing level determined according to the first two criteria (see **Methods**; **Fig. 2D**). The rationale behind the use of the three features for predicting functional back-splicing (**Fig. 1**) implies that applying stricter thresholds of these features will likely improve the predictive power of COL. When we set 4× to 20× the mean Cook’s distance as the cutoff, we indeed observed that the predictive power of COL rose with the threshold (**Fig. 3A**). For example, when the cutoff of Cook’s distance is 20×, 62.5% of the identified back-splicing is shared, significantly higher than the 31.4% under the threshold of 4× (**Fig. 3A**). Furthermore, we predict that considering motifs conserved between human and mouse, which diverged ∼90 mya [33], should make COL perform better than considering motifs conserved between human and macaque, which diverged 23 mya [34]. This prediction is indeed correct (**Fig. 3B**). In addition, considering motifs shared by all three species in COL outperforms considering those shared by only two species (**Fig. 3B**). Because the back-splicing level cutoff is determined by the other two features in COL (see **Methods**), we did not examine its effect separately. Together, the above analyses suggest that increasing the thresholds of outlier and motif conservation can improve the predictive power of COL.

**Fig. 3.**
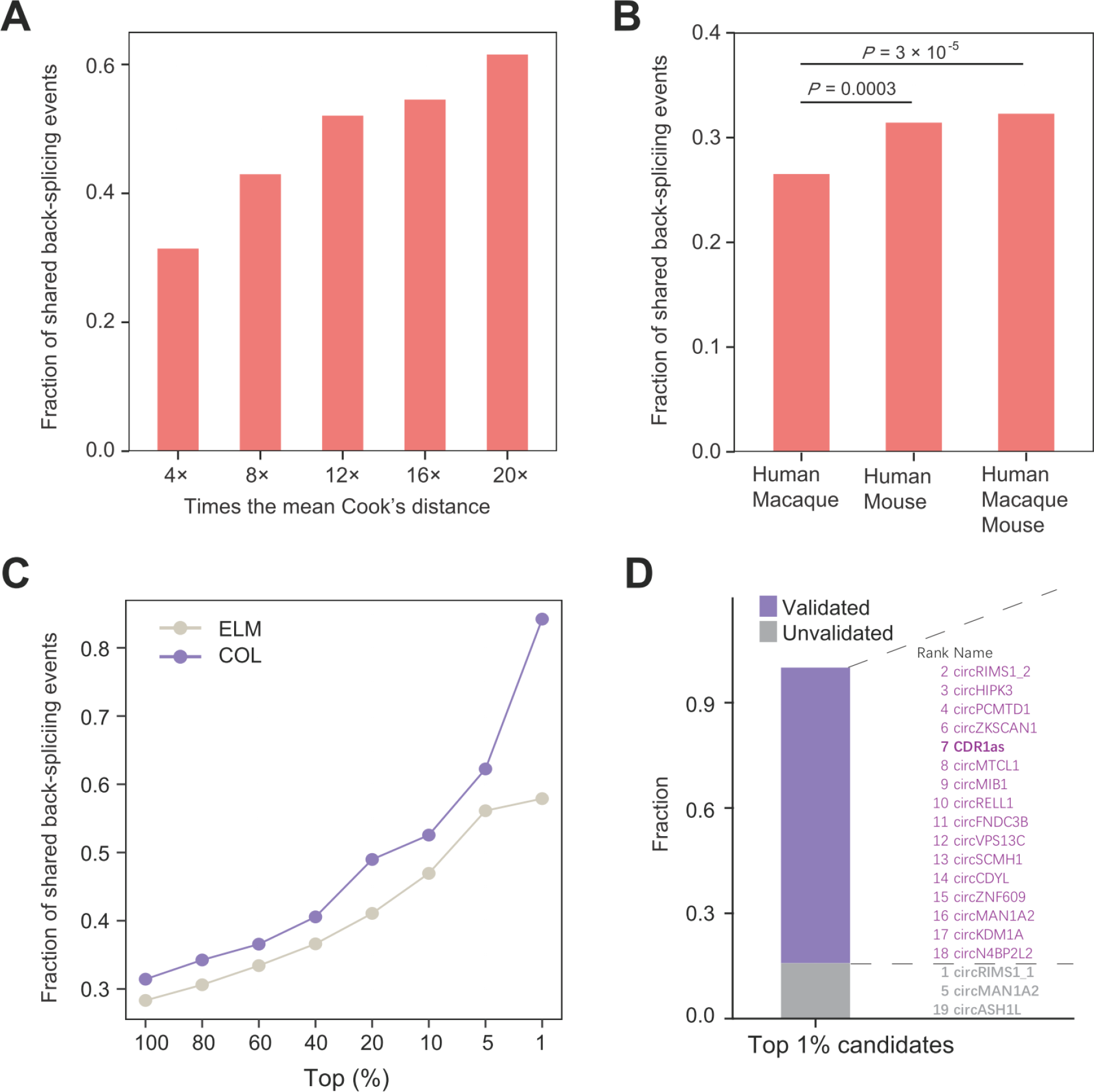
Performance of COL. (**A**) Fraction of back-splicing events identified by COL (under different cutoffs of Cook’s distance) that are shared. (**B**) Fraction of back-splicing events identified by COL (using motifs with different levels of conservation) that are shared. *P*-value is from Chi-square test. (**C**) Fraction of back-splicing events in different top proportions of candidates identified by COL or ELM that are shared. (**D**) Fraction of experimentally validated back-splicing events in the top 1% of the back-splicing candidates identified by COL.

In COL, potentially functional back-splicing events are prioritized by RRA. Highly ranked back-splicing events are presumably more likely to be functional. To validate this prediction, we selected different proportions (from top 100% to top 1%) of the ranked candidates and calculated the fraction of shared back-splicing (**Fig. 3C**). We found the fraction of shared back-splicing rises with the rank priority of candidates (**Fig. 3C**). Remarkably, this fraction reaches 84.2% (i.e., 16 of 19 candidates) for the top 1% of candidates. Therefore, in addition to raising the thresholds of the features, selecting top-ranked candidates can significantly improve the predictive power of COL.

To further assess the performance of COL, we searched the literature for experimental evidence for the functionality of the top 1% of the back-splicing events identified by COL. We found such experimental evidence for 16 of the top 19 candidate back-splicing events (**Fig. 3D; Data S1**). For example, the best known circRNA, CDR1as [7, 8], is ranked the seventh in our list. Even for the three back-splicing events without explicit experimental evidence for functionality, two may be functional (**Data S1**). Specifically, the top ranked circRIMS1-1 shares the host gene and donor site with the functional circRIMS1-2 [35, 36] and circMAN1A2 was reported to be strongly upregulated in cervical adenocarcinoma [37].

### Comparing the performance between COL and other methods

We next compared the performance of COL with those of other methods. First, we randomly picked the same number (i.e., 1,960) of back-splicing events as identified by COL and examined the number of shared back-splicing events. This was repeated 1000 times to acquire the mean number. We found on average 54.7 shared back-splicing events, significantly fewer than the 616 identified by COL (*P* < 0.001; **Fig. 4A**).

**Fig. 4.**
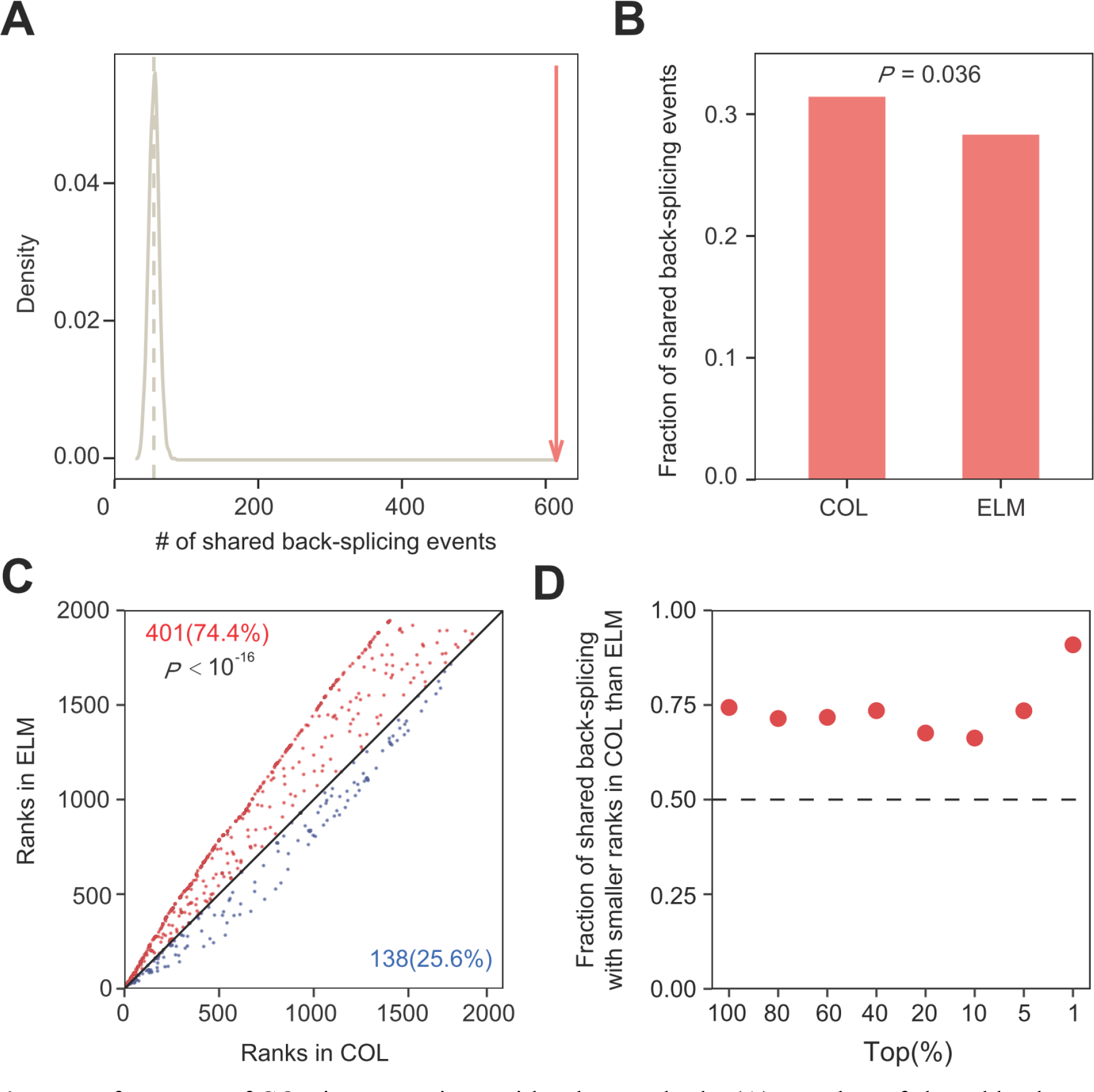
Performance of COL in comparison with other methods. (**A**) Number of shared back- splicing events identified by COL (red) or randomization (grey distribution). (**B**) Fraction of back-splicing events identified by COL or ELM that are shared. *P*-value is from a chi-squared test. (**C**) Ranks of shared back-splicing events. Shared back-splicing events identified by COL are ranked by RRA whereas those identified by ELM are ranked by the back-splicing level. Dots above and below the diagonal are respectively colored in red and blue, with their numbers and fractions indicated. The binomial *P*-value is presented. (**D**) Fraction of shared back-splicing events with smaller ranks in COL than in ELM in various top proportions of candidates identified by both methods.

We then compared COL with ELM, which is the only existing method that does not require more information than what COL uses. As mentioned, ELM detects putatively functional circRNAs by their high expressions. Because the back-splicing level, which is equivalent to the resulting circRNA expression level, is one of the three considerations of COL, COL should outperform ELM. We indeed found that the fraction of shared back-splicing events identified by COL is significantly higher than that identified by ELM (*P* < 0.04, chi-squared test; **Fig. 4B**).

This trend remains regardless of the specific top proportion of candidates considered (**Fig. 3C**). In particular, for the top 1% of the candidates, the fraction of shared back-splicing identified by COL is 45.4% higher than that identified by ELM (0.579) (**Fig. 3C**).

To further compare COL with ELM, we examined the rankings of the shared back-splicing candidates identified by the two methods (a smaller ranking indicates a higher probability of being functional). Of the 542 shared back-splicing events identified by both methods, 74% have smaller rankings in COL than ELM, significantly more than the random expectation of 50% (*P* < 10^-16^, binomial test; **Fig. 4C**). Qualitatively similar results were observed in different top proportions of candidates (**Fig. 4D**), suggesting that COL performs better than ELM in rankings of putatively functional circRNAs identified.

## DISCUSSION

Identifying the small fraction of millions of circRNAs that are potentially functional is a difficult task. In this study, we used back-splicing that is shared among human, macaque, and mouse as a proxy of functional back-splicing. We first demonstrated three features of functional back-splicing, and then developed the computational pipeline COL and its R package by considering and integrating the three features. We then showed that increasing the thresholds of the three features and prioritizing highly ranked candidates can significantly improve the predictive power of COL (**Fig. 3**). In particular, the fraction of shared or functional back-splicing can reach 84.2% among the top 1% of candidates identified by COL. Nonetheless, it should be noted that while these treatments increase the fraction of functional back-splicing among the predicted candidates (i.e., reducing the false positive rate), they also filter out functional back-splicing and result in a lower true positive rate; this may be less of a concern because at this stage the main task is to predict functional back-splicing precisely rather than comprehensively.

Compared with other methods for discovering functional circRNAs, COL has its own advantages. Unlike DEM that requires case and control samples, COL can be applied to a single sample. Different from CM that depends on back-splicing data from multiple species, COL needs back-splicing data from only one species (plus genome sequences from other species).

Although ELM has the same advantage of depending on a single sample, it is substantially inferior to COL in performance (**Fig. 3**).

The three features considered in COL have different predictive powers in identifying shared back-splicing. In particular, splicing motif conservation seems to be the least powerful feature (1.5-fold enrichment of shared back-splicing relative to the baseline), but it can substantially improve the prediction when used together with another feature. For example, the combination of regression outliers and splicing motif conservation achieves a 9.86-fold enrichment (**Fig. 2B**), which is even higher than the sum of the individual enrichments they bring (7.2 + 1.5 = 8.7; **Fig. 2A**).

Recently, a CRISPR-Cas13-mediated circRNA screen (CDCscreen) found 63, 62, and 67 putatively functional back-splicing events in Hela, 293FT, and HT29 cell lines, respectively [29], from which 9 candidates in each cell line were knocked down and 8, 7, 8 of them were confirmed by the phenotype of reduced cell proliferation (referred to as KD-validated).

Although the functionality of these circRNAs was based on experiments, we did not use them to assess COL because of several shortcomings. First, this dataset was based on the test of a very small fraction (762 of 12,338) of back-splicing events detected in the cell lines. Second, it was limited to the function of cell proliferation. Third, it was biased toward ELM because the 762 candidates were selected according to the back-splicing level. Nevertheless, COL can find most of the KD-validated back-splicing events (i.e., 7 of 8 in Hela, 4 of 7 in 293FT, and 4 of 8 in HT29). Several other high-throughput approaches for screening functional circRNAs [38, 39] have similar drawbacks so were likewise not considered here.

Apart from the three features considered in COL, functional back-splicing likely has other properties. For example, back-splicing is regulated by *trans*-factors and *cis*-elements. Because *trans*-factors are the same for all back-splicing events in a cell whereas *cis*-elements are variable, the latter may be able to differentiate functional from non-functional back-splicing.

Although COL already considers the conservation of splicing motifs, which are *cis*-elements, other *cis*-elements regulating back-splicing may be integrated in the pipeline in the future. For instance, complementary sequences in the intron adjacent to the splicing junction, especially Alu sequences, are key regulators mediating back-splicing [40–43]. However, due to the complexity of complementary sequences [40], it is not easy to evaluate their conservation among species.

Therefore, we decided not to consider it at this stage.

Whether a back-splicing event is functional may also be inferred from the properties of its resultant circRNA. Binding to miRNAs or proteins is presumably a main function of functional circRNAs [7, 8]. Thus, if the miRNA- or protein-binding motif of a circRNA is evolutionarily conserved, the circRNA and the corresponding back-splicing are likely functional. Indeed, most of the experimentally confirmed functional circRNAs from the top 19 back-splicing events identified by COL function through miRNA- and/or protein-binding (see references in **Data S1**). In addition, some programs and webservers use binding information to predict circRNA functionality [44–46]. However, due to frequent alternative splicing inside circRNAs [47], the sequence and length of the circRNAs resulting from the same back-splicing are variable. The short reads from RNA-seq capture only back-spliced junctions and do not provide the whole circRNA sequence, hampering the consideration of binding motif conservation in COL. Because circRNAs can function through its protein products [11], translational activity would be another indicator of circRNA functionality. However, proteomic data are currently lacking in most studies, so the translation information is unavailable. These additional features may be integrated into COL when the relevant techniques become better or relevant data become available. However, we must caution that demonstrating a biochemical activity of a molecule (*e.g.*, binding to a protein or subject to translation) does not mean that the molecule has a biological function and the ultimate proof of the functionality of a molecule is that it is subject to selection [48].

## METHODS

### RNA-seq datasets

Our study requires datasets of both back-splicing and linear-splicing from the same sample. RNA-seq libraries enriched by oligo(dT) or treated by RNase R are inappropriate because the former filters out RNAs produced by back-splicing while the latter removes RNAs generated by linear-splicing. Thus, we used RNA-seq data generated from total RNA samples treated with only the RiboMinus Kit because it removes rRNAs but retains both back-splicing and linear-splicing products. Specifically, the RiboMinus RNA-seq datasets from Ji P, et al. [3] downloaded from NGDC (https://ngdc.cncb.ac.cn/<u>; ID: PRJCA000751</u>) were used in our study. They include data from multiple tissues of human, macaque, and mouse. In addition, we used the RiboMinus RNA-seq datasets of HT29, 293FT, and Hela cell lines reported by Li S, et al. [29] and downloaded from GEO (https://www.ncbi.nlm.nih.gov/geo/; ID: GSE149691). All of the RNA-seq datasets were processed by FastQC for quality control, to remove adaptors and low-quality reads by Trimmomatic, and to filter out rRNAs through Bowtie alignment.

### Linear-splicing and back-splicing

For the datasets from Ji P, et al. [3], linear-splicing and back-splicing junctions were retrieved following our previous study [20]. Briefly, we used the genome assemblies of GRCh38 for human, Mmul 8.0.1 for macaque, and GRCm38 for mouse, all downloaded with gene annotations from Ensembl release 89. Linear-splicing was inferred by Tophat2 [49] using default parameters, including annotated and newly identified splicing. Back-splicing was identified by CIRCexplorer2 [40], which was then merged with newly identified back-splicing by CIRI2 [50, 51].

Li S, et al. [29] analyzed their data using GRCh37. To be consistent with their results, we analyzed their data using the same human genome reference (Ensemble release 75). Specifically, linear-splicing was identified by STAR (v2.4.2) using default parameters. We followed the original authors to identify back-splicing. That is, HISAT2 was used to map clean reads to the reference genome; the unmapped reads were collected by Samtools and further mapped to the reference genome by TopHat-Fusion. Finally, CIRCexploere2 was used for parsing back-splicing from the mapping results of TopHat-Fusion.

### Splicing amount, back-splicing rate, and back-splicing level

A splicing event has an acceptor site and a donor site. The total splicing amount at a splicing junction is the total number of linear- and back-splicing reads of its acceptor and donor sites. The back-splicing rate is twice the number of back-spliced reads divided by the total splicing amount at the back-splicing junction (**Fig. S3**). The relative back-splicing level at a back-spliced junction is the number of back-spliced reads at the junction per million total back-spliced reads in the sample (BSRPM).

### Cook’s distance

Cook’s distance is used in regression analysis to find influential outliers [52]. The formula for calculating Cook’s distance of data point *i* is 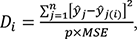, where *ŷ*_*j*_ is the *j*th data point’s fitted response value, *ŷ*_*j*_(*i*) is the *j*th data point’s fitted response value when data point *i* is removed, *n* is the number of data points, *MSE* is the mean squared error of the regression model, and *p* is the number of coefficients in the regression model. Four times the mean Cook’s distance of all data points was used as the cutoff to call outlies in our study. The commonly used cutoff advised by statistical tools R [53] and MATLAB [54] is three times. We used a stricter cutoff to guard against false positives.

### Ordering of back-splicing events

Back-splicing events were ordered using several different methods, including Average, Robust Rank Aggregation (RRA) [32], and Stuart [55]. The Average method simply takes the mean of the ranks of a target from multiple samples. RRA and Stuart are both based on order statistics in rank aggregation, but the former is more robust than the latter to noise [32]. All orderings were executed by the R package RobustRankAggreg. In addition, back-splicing events were ordered by their Cook’s distances or back-splicing levels.

### Splicing motif conservation and shared back-splicing

Back-splicing motifs, same as linear-splicing motifs, are GU adjacent to the donor site and AG adjacent to the acceptor site. We used the LiftOver (http://genome.ucsc.edu/cgi-bin/hgLiftOver) to identify orthologous genomic loci across species. If the nucleotides of a back-splicing motif in one species are the same as the nucleotides of the orthologous locus in another species, this motif is conserved; otherwise, it is unconserved.

If a back-splicing event in one species is also observed at the orthologous locus of another species, the back-splicing event is considered shared between the two species. In this study, shared back-splicing refers to back-splicing events shared among human, macaque, and mouse.

### Implementation and availability

The implementation of COL is composed of four sequential steps (**Fig. 2D**): (1) It first regresses between the back-splicing rate and splicing amount across splicing events and calculates a Cook’s distance for each back-splicing event. Then, back-splicing events that are above the regression line and have a Cook’s distance exceeding four times (or other times) the mean Cook’s distance of all back-splicing events (i.e., the outliers) are chosen as the potentially functional candidates. (2) Subsequently, the genomic motifs (i.e., GT-AG) of these candidate back-splicing events are examined for conservation across different species (i.e., human versus mouse in this study); those that are unconserved are eliminated. (3) Next, the same number of back-splicing events as that from step (2) are selected based on the expression level rank from high to low. These events are integrated with the result of step (2) through the intersection of Venn diagram to acquire the final list of potentially functional back-splicing events. (4) Finally, these candidates are prioritized by RRA ranking method based on the Cook’s distances and back-splicing levels.

The R package of COL is freely available at https://github.com/XuLabSJTU/COL.

### Data visualization and statistical analyses

R (v3.5.1) was used in statistical analyses and data visualization. Figures were generated using R packages, including ggplot2 and its dependencies.

## ACKNOWLEDGEMENTS

We thank Dr. Wei Xue for his help in processing the data from Li et al (2021). This study was supported by National Natural Science Foundation of China (32270704 and 32100518), National Science and Technology Innovation 2030 Major Projects for ‘Brain Science and Brain-Inspired Research’ (2022ZD0214400), Medical-Engineering Crossover Fund of Shanghai Jiao Tong University (YG2022QN084), and the U.S. National Institutes of Health (R35GM139484 to J.Z.).

## AUTHOR CONTRIBUTIONS

C.X. and J.Z. conceived the study and wrote the paper. Z.L., C.X., F.Z., T.Z performed the study and analyzed the data, B.S. developed the R package of COL.

---

**Fig. S1.**
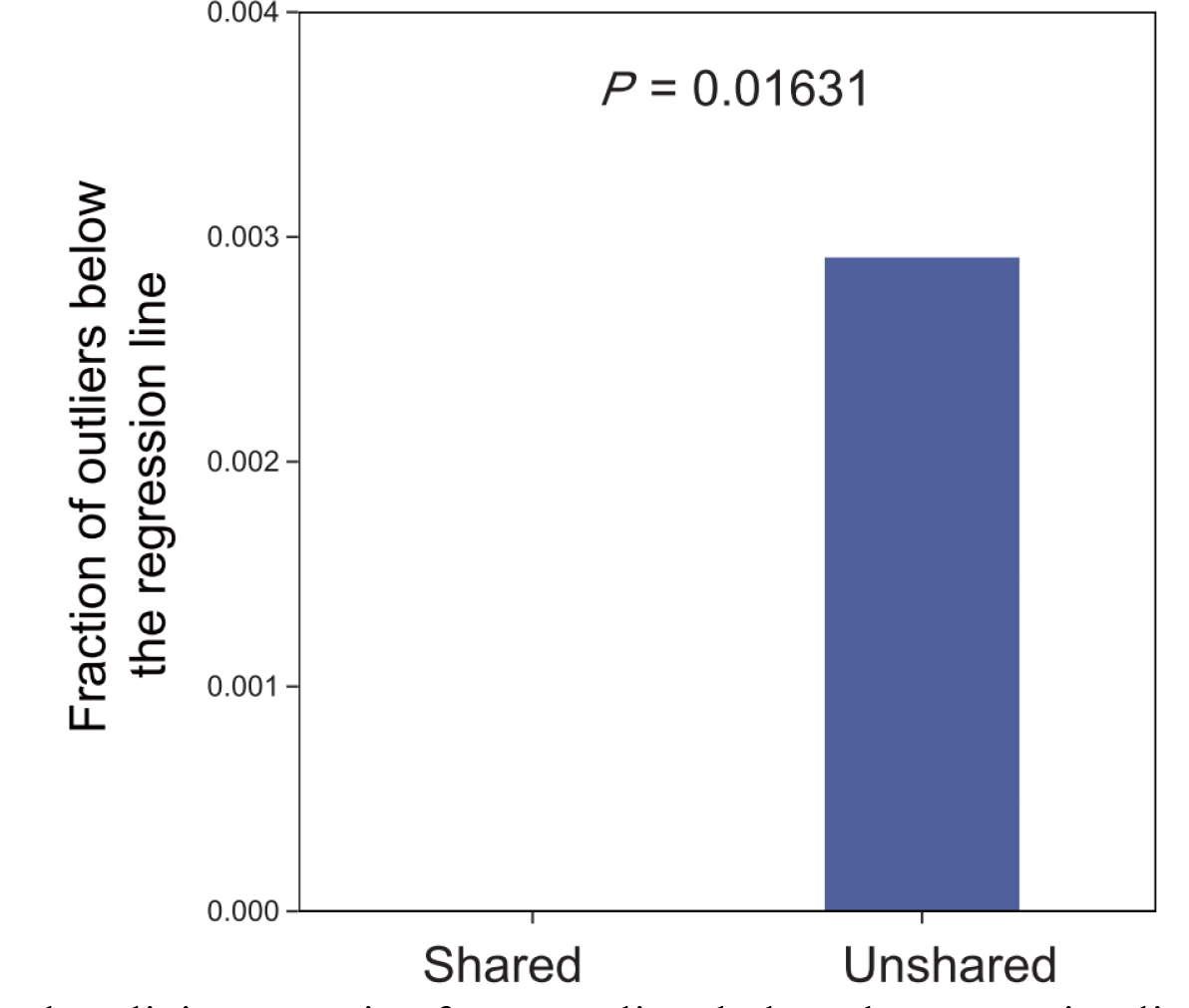
Shared back-splicing contains fewer outliers below the regression line than does unshared back-splicing.

**Fig. S2.**
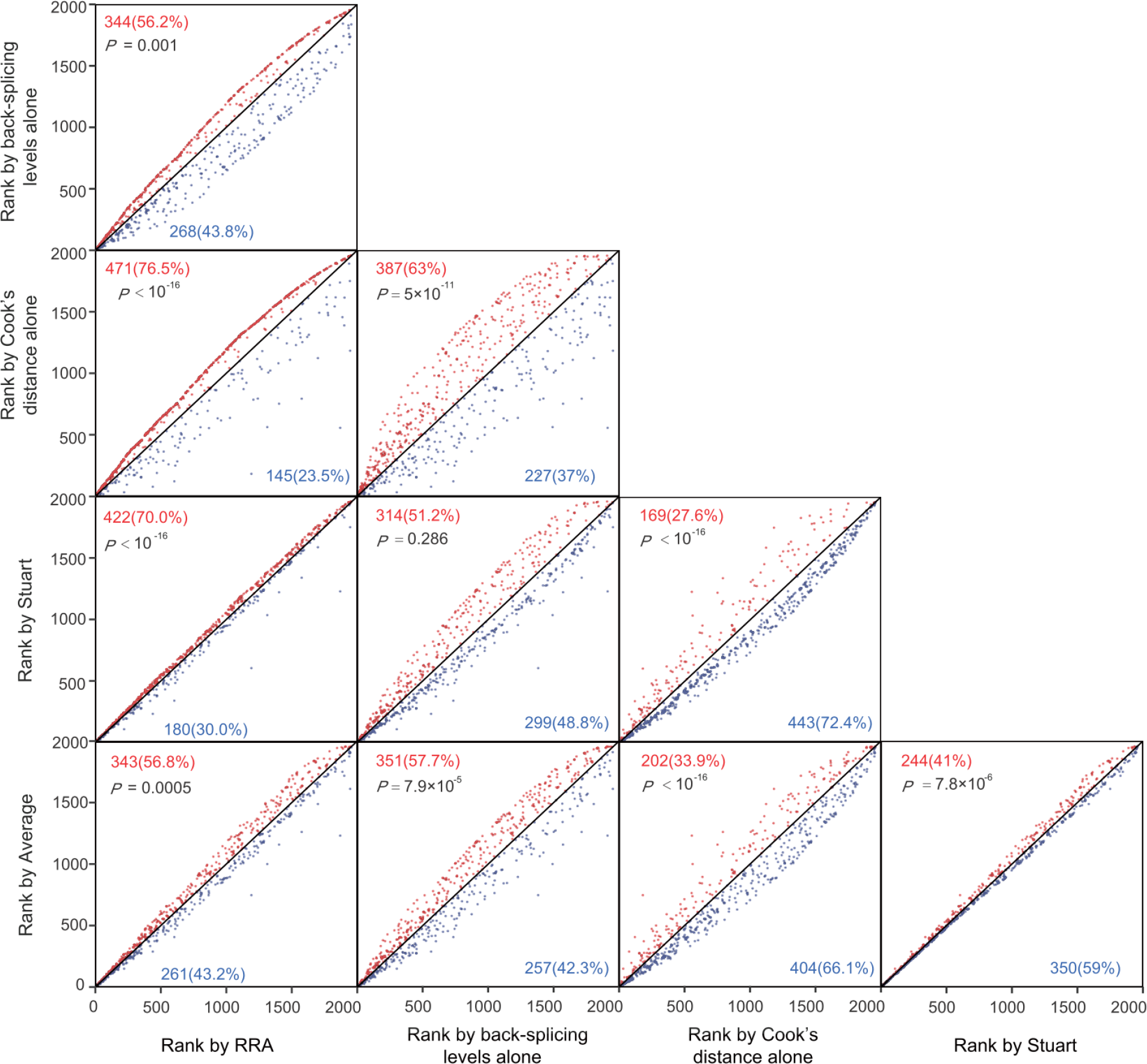
Pairwise comparison between ranking methods. Each dot represents a back-splicing event. Dots above and below the diagonal are respectively colored in red and blue, with their numbers and fractions indicated. *P*-values are from binomial tests.

**Fig. S3.**
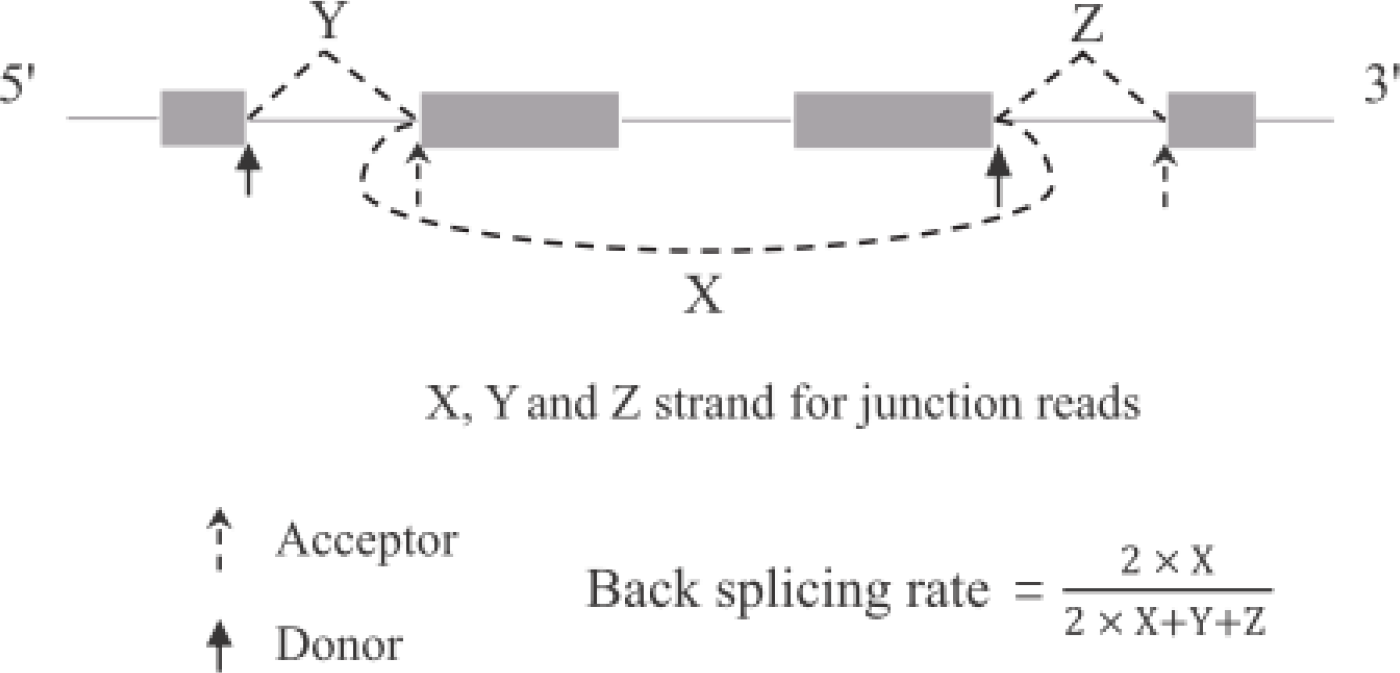
Diagram explaining the calculation of the back-splicing rate at a back-splicing junction.

**Data S1.** Back-splicing events identified by COL. The top 1% of back-splicing events (i.e., top 19) are yellow highlighted and manually checked for experimental evidence in the literature for functionality.

